# Continuous Biomarker Assessment by Exhaustive Survival Analysis

**DOI:** 10.1101/208660

**Authors:** Dominic A. Pearce, Ajit J. Nirmal, Tom C. Freeman, Andrew H. Sims

## Abstract

Publicly available high-throughput molecular data can enable biomarker identification and evaluation in a meta-analysis. However, a continuous biomarker’s underlying distribution and/or potential confounding factors associated with outcome will inevitably vary between cohorts and is often ignored. The *survivALL* R package (https://CRAN.R-project.org/package=survivALL) allows researchers to generate visual and numerical comparisons of all possible points-of-separation, enabling quantitative biomarkers to be reliably evaluated within and across datasets, independent of compositional variation. Here, we demonstrate *survivALL’s* ability to robustly and reproducibly determine an applicable level of gene expression for patient prognostic classification, in datasets of similar and dissimilar compositions. We believe *survivALL* represents a significant improvement over existing methodologies in stratifying patients and determining quantitative biomarker(s) cut-points for public and novel datasets.

## INTRODUCTION

Biomarker performance is traditionally assessed by estimating survival benefit over time, often summarized by Kaplan-Meier plot^1,2^. Whilst qualitative biomarkers can easily stratify cohorts into two or more groups, the question of how to divide a cohort with a quantitative measure is considerably more challenging. Multiple subgroups may exist within a given patient population, but a need for simple and tractable clinical decisions (i.e. treat or don’t treat) have often encouraged division into two classes. The most obvious, common, yet arbitrary approach for this is to divide a cohort into two equal sized groups at the median level. However, this median-split approach ignores a marker’s distribution and any potential confounding factors relating to the composition of the dataset, clinical or otherwise. This heterogeneity makes assessing the robustness of a biomarker across datasets problematic, with median dichotomisation unlikely to produce reliable results due to random sampling differences. In short, a significant median separation in cohort A is unlikely to translate to cohort B. Given the public availability of high-throughput molecular profiling datasets providing outcome data, the opportunity to evaluate biomarkers *in silico,* as part of extended meta-analyses, is in principle becoming easier. In practice however, datasets derived from cohorts of primary patient tissue are molecularly heterogeneous in their composition^3^, are likely to exhibit varying proportions of multiple clinical factors associated with outcome (e.g. node and receptor statuses, grade, molecular subtypes and age). It is therefore clear that this variation must be accounted for if a biomarker is to be successfully applied and validated in a meta-analysis.

Building upon previous best-of-split methods^4^–^6^ we have developed *survivALL* (https://CRAN.Rproject.org/package=survivALL) an R package implementation to exhaustively calculate and visualise hazard ratios (HRs) for all possible points-of-separation and assess the association between a continuous measure and survival. As open-source software, *survivALL* allows for researchers to reproducibly perform exhaustive survival analysis using public or independent datasets, in a highly transparent, automatable and extensible manner within the wider R package landscape. Complimenting *survivALL* is a companion web-based app – *survivAPP* (pearcedom.shinyapps.io/survivapp/) – allowing non-programmatic, drag-and-drop *survivALL* use. We believe that *survivALL* allows researchers to perform exhaustive survival analysis relevant to the current state of –omics research.

## MATERIALS AND METHODS

*survivALL* computes hazard ratio statistics for every point-of-separation possible, allowing the magnitude, and frequency of, significant cut-points to be identified (Video S1)x. To more robustly determine significance, a non-parametric and dataset-specific bootstrap is applied, defining reliable confidence intervals. Bootstrapping is performed as a 10,000-fold repeated calculation of HRs for random sample orderings, producing a distribution of expected/random HRs for each individual point-of-separation, to which observed true biological HRs are compared and significance calculated. Dataset-specific best points-of-separation were optimised by maximising desirability of combined p-value and absolute HR magnitude^7^.

All analysis was performed using the R statistical environment using CRAN and Bioconductor hosted packages^8^–^10^. *survivALL* p-values were calculated using the bootstrapping procedure described above, whilst median approach p-values were calculated using the Cox-proportional hazards model^1^. To avoid overcorrection of the multiple comparisons inherent in *survivALL*, all *survivALL* analysis is preceded by a single ancillary test of significance (Cox-proportional hazards model), performed using the biomarker as a continuous variable. Code and session information necessary to reproduce the analysis is included as supplementary material.

## RESULTS

To illustrate the value of the *survivALL* package, we considered over-expression of human epidermal growth factor 2 (HER2/*ERBB2*), a well-established biomarker associated with poor prognosis in invasive breast cancer^11,12^, in the largest single breast cancer gene expression dataset – METABRIC^13^ (*n=1971* samples with complete disease specific survival information). Importantly and unusually, METABRIC is split into two equally sized and composition-matched subsets, allowing for independent discovery and validation. The *survivALL* plotALL() function enables visualisation of hazard ratios for all possible separations (n-1) of ordered *ERBB2* expression for the METABRIC discovery cohort (*n=980*) (Figure 1A). It establishes 283/980 points-of-separation as significant, confirming the established expectation that increased *ERBB2* expression is associated with poor prognosis. The span and location of these points are consistent with epidemiological evidence demonstrating that across the population ∼20% of breast cancers overexpress HER2^14^. Contrasted in Figure 1B are two Kaplan Meier plots resulting from using the median (p=0.48) and the data-driven most significant point-of-separation (p=1.25×10^−11^) for this dataset.

**Figure 1:**
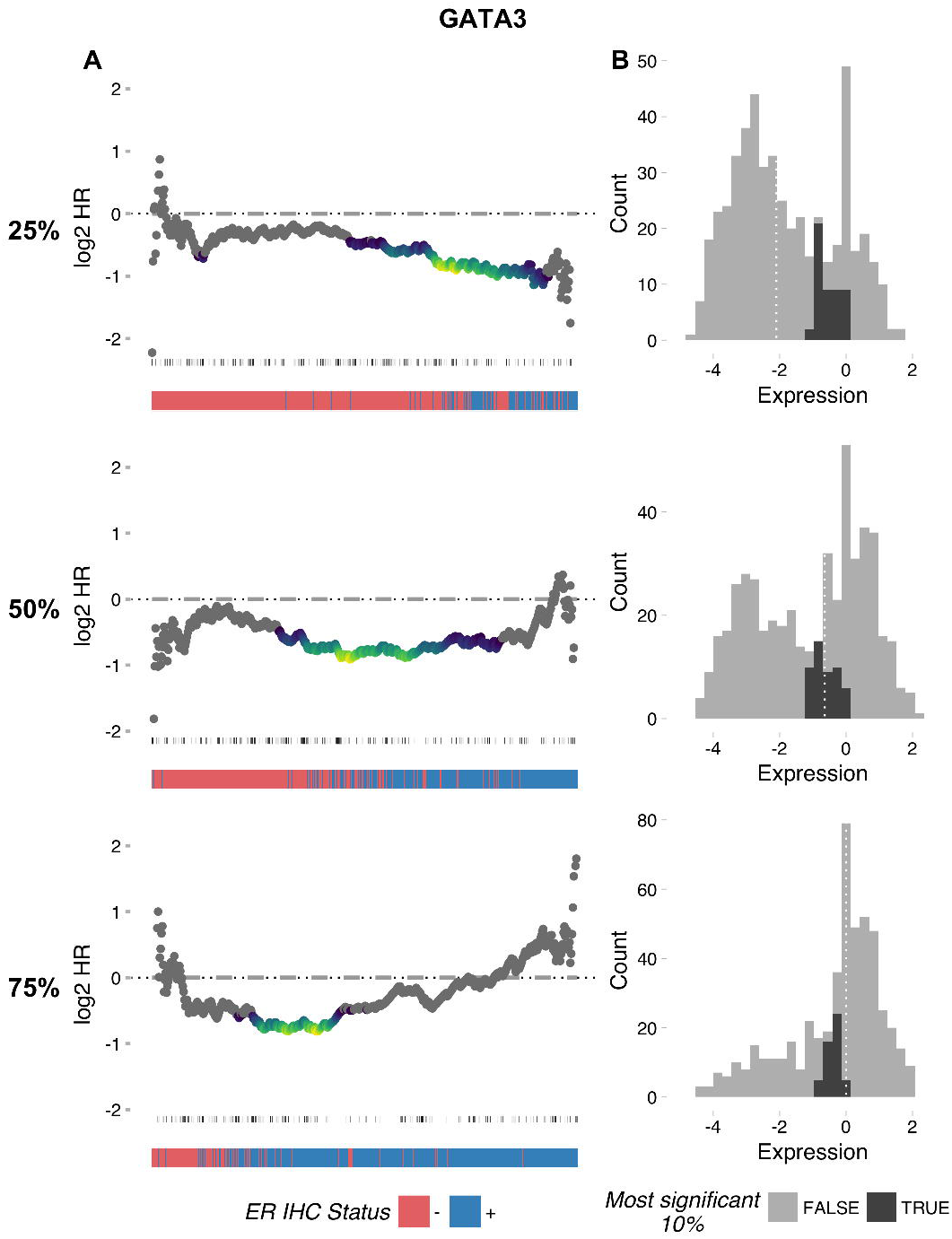
A) The plotALL function allows hazard ratios (y-axis) and bootstrap significance (p<0.05 colour scale; p 0.05 grey) for all possible quantitative biomarker cut-points to be examined. Patients are ordered by increasing expression of the quantitative marker – in this example, *ERBB2* expression. Events (distant metastasis-free survival) are indicated as vertical lines below the plot, with darker colours indicating chronologically earlier events. Kaplan-Meier plots for *ERBB2* stratified using median (n = 490 vs. 490) and survivALL (n = 924 vs. 56) approaches are shown, comparing patients with expression lower (grey) or higher (black) than the points-of-separation determined in A. B) Comparison of *survivALL* and median approaches in predicting optimal cohort separation for *ERBB2*. The expression values associated with both median and dataset-specific best points-of-separation derived from the discovery set in 1A is applied to the validation dataset. Error (median = orange, *survivALL* = blue) is calculated as the number of patient who would be incorrectly assigned using these predictions, in relation to the validation cohorts own best separation. This process is repeated in 1C for all genes. *survivALL* significantly (p=2.2×10^−16^) outperforms a median approach in terms of predictive accuracy.

In considering the issue of dataset composition, the highly similar METABRIC subsets enabled us to first evaluate whether *survivALL* could derive and validate the point-of-separation which most clearly distinguishes (lowest p-value) between good and poor prognosis groups from one cohort to the other (Figure 1B). For all 19,628 genes, *survivALL* was used to calculate the point-of-separation with the lowest p-value in the discovery cohort and the gene expression value at this point was used to divide the validation cohort (*n=991*). Prediction accuracy was measured as the number of patients incorrectly classified compared to the validation cohort’s own dataset-specific most significant point-of-separation. This same measure of accuracy was additionally calculated using a median approach for comparison (Figure 1B & 1C).

As expected, *survivALL* significantly outperforms a median approach (p=2.2×10^−16^, Kolmogorov-Smirnov test), demonstrating *survivALL*’s applicability in robustly determining prognostic stratification in two suitably relatable datasets, with no *a priori* knowledge of a gene’s population level distribution.

However, real-world datasets are rarely compositionally matched in the way that the METABRIC discovery and validation cohorts are, and it remained to determine if *survivALL* could perform this applied prognostic stratification in more dissimilar and noisier datasets. To test this, we simulated semi-random sub-samplings of the entire METABRIC dataset for three pre-defined proportions of estrogen receptor (ER) positivity – 25, 50 & 75%. Using these dramatically variable compositions, we determined the extent to which *survivALL* was able to track these differences in terms of the datasets-specific best point-of-separation of *GATA3*, a mediator of ER binding^15^ (Figure 2A). It was evident that as the proportion of ER+ and ER-samples shifted, the *survivALL* plots in turn shifted in response, with the most significant point-of-separation consistently falling at the division between ER+ and ER-samples. Importantly, though these plots changed, the corresponding level of expression that defined the ER+/-boundary remained consistent (Figure 2B), indicating *survivALL’s* ability to robustly determine a reliable level of GATA3 expression to stratify our patient cohorts, even in highly compositionally dissimilar datasets. We repeated this analysis for 5 publically available, ER-positive and tamoxifen treated datasets (GSE2990, GSE6532, GSE9195, GSE12093, GSE17705), for grade and AURKA expression, demonstrating similarly consistent results between datasets (Figure S1).

**Figure 2:**
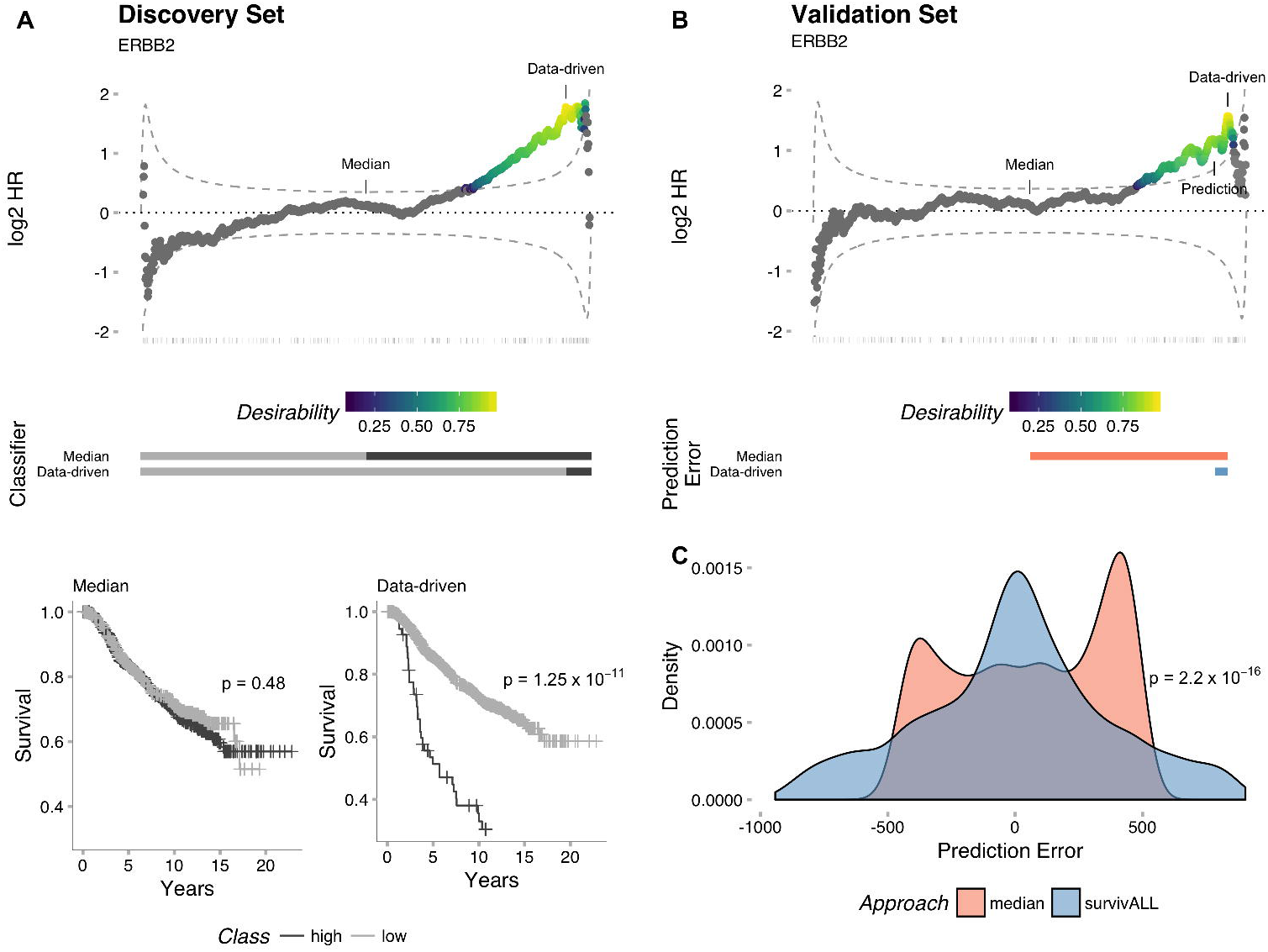
A) Analysis of compositionally *un*matched METABRIC subsets. Three equally sized subsets (n=500) with variable proportions of ER+ samples (25, 50 and 75%) demonstrate *survivALL*’s ability to track the ER+/ER-(blue/red) boundary for *GATA3* expression, with the most significant point-of-separation (low-high significance = blue-yellow gradient) existing at this boundary for each subset. B) Whilst the HR distribution is evident to change with ER+ proportion, the 10% most significant points-of-separation consistently related to the same level of *GATA3* expression. For comparison, the median level of expression (dashed white line) for each subset is also shown.

Finally, whilst significant association of a biomarker can be determined in a meta-analysis using expression as a continuous variable, this does not reveal the direction of that association, i.e. good or poor prognosis. Beyond the added information revealed therein, there additionally remains the possibility that a gene determined as significantly associated with prognosis in more than one dataset may in fact demonstrate variable, or even opposite, directions of association (Figure S2). For the METABRIC discovery and validation cohorts this was evident to occur for ten genes, several of which (*ACY3*^16^, *LRRK2*^17^, *NUPR1*^18^ & *UGT1A7* ^19^) have been previously associated with cancer risk. This therefore represents a small but real danger that must be considered in meta-analysis.

## DISCUSSION

In this study we have highlighted the issue of assessing quantitative biomarkers for survival analysis, offering *survivALL* and *survivAPP* (Video S2) as tools to evaluate and overcome these challenges. We have attempted to illustrate situations relevant and common to researchers in the current environment of publically available large datasets, using well established but likely compositionally distorted examples. As an R package, *survivALL* allows greater resolution, transparency and flexibility compared to other online best-of-split tools, applicable to any public or proprietary dataset and usable in larger-scale automated data operations. Researchers are increasingly using approaches such as KMplotter^20^, either to median-split or highlight the dataset-specific best point-of-separation in a single or combined dataset. This approach presents a number of potential problems, most notably the restriction of what data is available to be analysed – either alone or in combination as a meta-analysis – and the exact methods used. Furthermore, whilst a prognostic association can be considered using a continuous marker as the classifier itself, this ignores a number of informative factors, such as the direction and magnitude of association, as well as the optimal value to allow patient stratification into treatment groups.

Importantly, whilst the examples presented here relate to individual genes in breast cancer microarray datasets, *survivALL* is readily extensible to other diseases, data types or any other quantitative measure, including signatures or scores based upon combinations of markers. Moreover, whilst this paper has largely considered dichotomisation, *survivALL* also allows for additional sub-populations with varying survivals to be visualised, including those related to clinical factors such as grade (Figure S1), as well as revealing potential confounding factors, producing multivariate analysis and demonstrating or uncovering an otherwise unknown factor when considering multiple datasets. Fundamentally, *survivALL* has been developed as open-source software, to flexibly integrate with other popular R packages, including the popular visualisation tool *ggplot2* for customisable and scalable output.

Most importantly, *survivALL* and *survivAPP* allows true biological effects and their relationship to survival to be revealed and reliably compared, within and between datasets, to move towards determining real-world clinically applicable biomarker cut-offs.

